# Pleiotropic functions of an antenna-specific odorant binding protein linking xenobiotic adaptation and olfaction in the Colorado potato beetle, *Leptinotarsa decemlineata*

**DOI:** 10.1101/2025.04.29.651160

**Authors:** James A. Abendroth, Timothy W. Moural, Casey Cruse, Jonathan A. Hernandez, Michael Wolfin, Tom C. Baker, Andrei Alyokhin, Fang Zhu

## Abstract

The Colorado potato beetle, *Leptinotarsa decemlineata*, is the primary defoliator of potatoes and is notorious for its robust capability to develop resistance to various insecticides used to control it. As the initial interface between the environment and the insect olfactory system, odorant binding proteins (OBPs) solubilize and transport hydrophobic odorant molecules from the sensillar lymph to olfactory receptors. Recently, evidence has suggested that OBPs may also sequester excess harmful molecules such as insecticides in the perireceptor space, preventing them from reaching vulnerable olfactory sensory neuronal dendrites. In this study, we identified an antenna-specific OBP (*LdecOBP33*) that is 2.5-fold higher expressed in a neonicotinoid resistant strain than the susceptible one. Competitive displacement fluorescence binding assays revealed that LdecOBP33 protein binds to a wide range of compounds including various plant volatiles and insecticides. Next, we used RNA interference to knock down *LdecOBP33* and combined with toxicological bioassays, behavioral, and electrophysiological assays to investigate functions of *LdecOBP33* in insecticide resistance and host location. We found that *LdecOBP33* silencing increased male beetles’ susceptibility to imidacloprid, suggesting its role in insecticide resistance. Additionally, *LdecOBP33* aids in host location through enhancing the detection of stress-induced potato plant volatiles, increasing the efficacy of CPB adult foraging. Taken together, our study is the first to functionally characterize an OBP in CPB linked to insecticide resistance and host location. Our findings provide insight into a key molecular factor involved in CPB’s response to environmental challenges, suggesting a potential link between insect adaptation to xenobiotics and olfactory processing.

## 1. Introduction

Insects are the most abundant group of metazoans on our planet, comprising a large portion of Earth’s biodiversity (Stork, 2018). A critical component to their success is the proper recognition of volatiles that act as chemical cues for potential hosts, conspecifics, and predators (Leal, 2013; Wei et al. 2021). Volatile detection primarily occurs in the antennae of terrestrial insects, which are finely tuned and extremely sensitive to perform this essential role (Baker et al., 1998; Pelosi et al., 2018). Odorant binding proteins (OBPs) are an important component of this system that act as the initial interface between the environment and the insect olfactory system (Abendroth et al., 2023). Typically, OBPs selectively bind and solubilize hydrophobic volatile molecules, protecting them from degradation before reaching the respective olfactory receptor (Brito et al., 2016; Leal, 2005). Recent evidence suggested that OBPs not only facilitate olfactory processing but also sequester excess harmful molecules in the perireceptor space, preventing them from reaching vulnerable olfactory sensory neuronal dendrites (Pelosi et al., 2018; Tricoire-Leignel et al., 2012; Whiteman and Pierce, 2008). These molecules include plant volatiles, pheromones, and pesticides, all of which can be harmful in excess to insect olfactory receptors (Abendroth et al., 2023; Cruse et al., 2023; Gong et al., 2009; Honson et al., 2003; Larter et al., 2016; Wang et al., 2020). OBPs may therefore be co-opted to both xenobiotic adaptation and olfaction (Abendroth et al., 2023).

Colorado potato beetle (CPB) (*Leptinotarsa decemlineata*) is a notorious agricultural pest, known for its robust capability to rapidly develop resistance to pesticides (Alyokhin et al., 2008; Grafius, 1997). In less than a century since the first synthetic pesticide was introduced, CPB has evolved resistance to 57 different active ingredients, ranking among the top 10 most pesticide-resistant arthropod pests in agriculture (Mota-Sanchez, 2024; Zhu et al., 2016a). It has been hypothesized that the CPB’s robust capacity to evolve pesticide resistance may be linked to its co-evolutionary history with solanaceous plants (Koirala BK et al., 2022; Zhu et al., 2016b). CPB are specialists of solanaceous plants, commonly known as the nightshade family. This family includes key agricultural crops such as potato, tomato, and eggplant, which contain high levels of toxic glycoalkaloids in various parts of the plant (Alyokhin et al., 2008; Chowanski et al., 2016; Lachman et al., 2001; Liu et al., 2021; Zhu et al., 2016b). Potato plants damaged by CPB can emit 7- to 10-fold increased amounts of herbivore induced plant volatiles, such as linalool, 1-hexanol, (E)-2-hexenal, and 2-phenylethanol, compared to the emissions of undamaged potato plants (Bolter et al., 1997; Sablon et al., 2013; Visser, 1979). These volatiles are monoterpenoids, C_6_-aldehydes, or alcohols that can have antimicrobial activities at high concentrations, potentially endangering adult CPB antennae (Bolter et al., 1997; Lachman et al., 2001; Sablon et al., 2013; Schutz et al., 1997b). Despite this challenge, CPB readily feeds on potato plant foliage at all developmental stages, indicating the presence of adaptive mechanisms that facilitate host plant utilization. While previous literature has well documented how insects adapt to toxic allelochemicals in their food (Heckel, 2014; Karageorgi et al., 2019; Li et al., 2007; Wang et al., 2018), little is known about how insects cope with host plant volatiles.

To bridge this knowledge gap, in this study we functionally characterized an antenna-specific OBP (*LdecOBP33*) that is highly expressed in adults, with particularly high expression in male antennae. Through purification of recombinant proteins, we assessed the binding affinity of LdOBP33 to various potato plant volatiles and classes of insecticides using competitive fluorescence displacement binding assays. After silencing *LdecOBP33* through feeding RNA interference (RNAi), we investigated its role in insecticide resistance and its impact on adult CPBs’ detection of potato plant volatiles. Recently, 59 OBPs were identified in the CPB genome, but none have been functionally characterized (Schoville et al., 2018). Here, we investigated a CPB OBP for the first time in relation to insecticide resistance and host location.

## 2. Materials and methods

### 2.1. Insects

Two different populations of CPBs were used for this study. The susceptible CPB was purchased from French Agricultural Research (Lamberton, MN, USA). This population of CPB was initially collected in 2003 from Long Island, NY, and was reared in laboratory conditions with no pesticide exposure for over fifteen years since then. The neonicotinoid-resistant CPB (Chen et al., 2014) was collected from the University of Maine Aroostook Research Farm (Presque Isle, ME, USA). Both CPB populations were subsequently reared in a Penn State facility greenhouse at 25 ± 1°C and a 16:8 L:D photoperiod regime. Beetles were fed on a constant supply of Red Norland potato plants in Insect and Butterfly habitat cages (Restcloud, Guangzhou, China). Eggs were collected from plants and stored in petri dishes maintained at 25 ± 1°C, 60% ± 5%, and under a 16:8 L:D photoperiod regime within a rearing room. Newly emerged first instar larvae fed upon fresh potato plant leaflets up to the third instar and then were transferred back to the greenhouse. After eclosion, males and females were separated into different habitat cages and were continuously supplied fresh potato plants.

### 2.2. In silico structural and phylogenetic analyses

*LdecOBP33* was the top candidate identified in our previous transcriptomic study of CPB antenna and exhibited 99% sequence similarity to XM_023165485 (Liu et al., 2021; Schoville et al., 2018). The signal peptide of LdecOBP33 was determined using SignalP 6.0 (https://services.healthtech.dtu.dk/services/SignalP-6.0/). Isoelectric point (pI) and molecular weight (mW) were estimated using ExPASy proteomic tools (http://web.expasy.org/compute_pi). Secondary and tertiary structure were predicted for LdecOBP33 using Alpha Fold 2 (Jumper et al., 2021; Mirdita et al., 2022).

A total of 219 OBP sequences from 5 different insect species (*Leptinotarsa decemlineata, Tribolium castaneum, Diabrotica virgifera virgifera, Monochamus alternatus*, and *Collaphellus bowringi*) were obtained from the NCBI database and prior published studies (Table S1). The multiple sequence alignment was performed using MUSCLE in MEGA 11 (Tamura et al., 2021) with the default parameters (Gap Open: -2.90, Gap Extend: 0.00, Hydrophobicity multiplier: 1.20). The maximum likelihood tree was inferred using MEGA 11, utilizing the LG model with gamma distributed with invariant sites (G + I), with a bootstrap replicate size of 1000 (Abendroth et al., 2023).

### 2.3. RNA extraction, cDNA synthesis, and qRT-PCR

Total RNA extraction was conducted using 3-50 CPB from each life stage: one-day old eggs, five-day old eggs, first to fourth instar larvae, pupae, and one week old adult males and adult females. Additionally, specific tissue was collected from one week old adult males and females, which included: antennae, head, legs, midgut, fat bodies, Malpighian tubules, and sex organs. Once collected, tissue was homogenized within TRIzol® reagent (Thermo Fisher Scientific, Waltham, MA, USA) according to the manufacturer’s instructions. RNA samples were then treated with Invitrogen Turbo™ DNAse (Thermo Fisher Scientific, Waltham, MA, USA) to eliminate genomic DNA contaminant. Purified RNA samples were then used to transcript to cDNA with M-MLV reverse transcriptase (Promega, Madison, WI, USA). NanoDrop One (Thermo Scientific, Madison, WI, USA) was used to measure the concentration of cDNAs. The qRT-PCR was carried out using a CFX96 Touch Deep Well Real-Time PCR Detection System (Bio-Rad Laboratories, Hercules, CA, USA). A 10 µL total reaction volume included 1 µL cDNA, 5 µL Forget-Me-Not™ EvaGreen qRT-PCR Master Mix (Avantor Inc., Radnor, PA, USA), 0.4 µL qRT-PCR primers (Table S2), and 3.6 µL ddH_2_O. A program comprising an initial incubation at 95°C for 3 minutes, 40 cycles at 95°C for 10 seconds, 55°C for 30 seconds, 95°C for 10 seconds, and lastly 65°C for 5 seconds was used for all reactions. Elongation factor 1α (EF1α) and ribosomal protein L4 (RPL4) were used as reference genes to normalize Ct (threshold cycle) values based on our prior studies (Moural et al., 2020; Zhu et al., 2011). Relative gene expression was determined with the 2^-ΔΔCt^ method (Livak and Schmittgen, 2001). Three biological replications and two to four technical replications were conducted independently.

### 2.4. RNA interference (RNAi)

The specific dsRNA of *LdecOBP33* was prepared using an Invitrogen MEGAscript™ T7 transcription kit (Thermo Fisher Scientific, Waltham, MA, USA) with primers listed in Table S2. The dsRNA of a green fluorescent protein (GFP) gene (ds*GFP*) was generated using a pET His6 GFP TEV LIC cloning vector plasmid (addgene#29663) as template. The reaction consisted of an initial incubation held at 37°C for 6 hours, then an extension step held at 75°C for 5 minutes, and finally an overnight annealing step at room temperature. Then the dsRNA was purified using an Invitrogen Turbo DNA-*free*™ Kit (Thermo Fisher Scientific, Waltham, MA, USA). The quality and length of dsRNAs was assessed through both agarose gel electrophoresis and a NanoDrop One Microvolume UV-Vis Spectrophotometer (Thermo Fisher Scientific, Waltham, MA, USA). The RNAi feeding procedure for dsRNA delivery to CPB adults was adapted from our previous studies (Moural et al., 2020; Zhu et al., 2011; Zhu et al. 2014). In brief, one-week-old adult male or female CPB were fed with 3 *µ*g dsRNA of *LdecOBP33* or ds*GFP* (control) on a potato leaf disc per beetle after 24 hours of starvation. The treatment lasted for three days, after which the beetles were fed untreated potato leaf discs for an additional two days. On the sixth day, CPB were collected for either knockdown efficiency evaluation by qRT-PCR, toxicology assays, or were starved for an additional 24 hours for use in behavioral and/or electroantennography (EAG) bioassays.

### 2.5. Heterologous expression and purification

A pET28a(+) plasmid with codon optimized full length *LdecOBP33* (XM_023165485) inserted between the NdeI and HindIII cut sites was ordered from GenScript™. Then the signal peptide and thrombin cut site were removed using a New England Biolabs Q5^®^ Site-Directed Mutagenesis Kit and the deletion primers listed in Table S1 to generate the final expression construct. Final plasmid construct was sequenced via Plasmidsaurus™ (Arcadia, CA, USA) to confirm construct accuracy. Then the plasmid was transformed into the Shuffle® T7 lysY competent *E. coli* expression strain (NEB, Ipswich, MA, USA) for expression. Cells containing pET28b-LdecOBP33 were incubated at 30°C overnight at 250 rpm, suspended in 50 mL of 2YT media (0.8 g tryptone, 0.5 g yeast extract, 0.25 g NaCl, 50 mL diH_2_O) and 100 *µ*g/mL of kanamycin antibiotic. After 16-18 hours, the culture was removed from the shaking incubator and was used to inoculate a 1.2 L expression culture of terrific broth (TB) media (24.0 g tryptone, 28.8 g yeast extract, 4.8 mL glycerol, 120 mL phosphate buffered saline (10x PBS), 1080 mL diH_2_O) supplemented with 100 *µ*g/mL of kanamycin. Following inoculation, the expression culture was incubated at 30°C at 250 rpm until an optical density (OD_600 nm_) value between 0.7-0.9 was reached, wherein the culture was then removed and chilled on ice for fifteen minutes, and then induced with 1 mM of Isopropyl ß-d-1-thiogalactopyranoside (IPTG).The culture was then subsequently incubated in a shaking incubator for 24 additional hours at 16°C and 250 rpm. Next, cells were pelted by centrifugation at 3.500 rpm and the pellet was stored at -20 °C until protein purification.

For LdecOBP33 purification the cell pellet was resuspended in a buffer solution containing 50 mM NaPi, 500 mM NaCl, 10 mM Imidazole, 3.3 mM NaN_3_, pH of 7.4, a Pierce Protease EDTA-free Inhibitor Tablet (Thermo Scientific), and 100 mM phenylmethylsulfonyl fluoride and lysed through sonication (Branson Digital Sonifier SFX 150) at 70% power for 30 seconds, a total of 3 to 5 times until a homogenous lysate was achieved. The lysate was then centrifuged at 18000 rcf for 15 min. The supernatant was transferred to a Kontes® Flex-Column® gravity column with a Ni-NTA resin bed and washed with ten column volumes (CV) lysis buffer and five CV of 50 mM NaPi, 500 mM NaCl, 40 mM Imidazole, 3.3 mM NaN3, pH of 7.4. Then LdecOBP33 protein was eluted with eight CV of 50 mM NaPi, 300 mM NaCl, 250 mM Imidazole, 3.3 mM NaN3, pH of 7.4 tube. This elution was then 100-fold buffer exchanged into 20 mM 2-morpholineethanesulfonic acid (MES), 1 mM ethylenediaminetetraacetic acid (EDTA), pH of 6.5, and injected onto a 5 mL HiScreen™Capto™ SP ImpRes chromatography column (Cytiva, Marlborough, MA, USA) connected to a Bio-Rad NGC. The column was washed until the 280 nm reading was at a constant steady baseline. Then a 20 CV gradient with 20 mM MES, 1 M NaCl, 1 mM EDTA, pH of 6.5 was used to elute LdecOBP33 protein. Purified LdecOBP33 was 100-fold buffer exchanged into 20 mM Bis-Tris, pH 6.5, and flash frozen with liquid nitrogen and stored at -80 °C until later use. Analysis of chromatography fractions and final purified protein was performed with sodium dodecyl sulfate polyacrylamide gel electrophoresis (SDS-PAGE), to assess the quality and quantity of LdecOBP33 protein. Concentration of purified LdecOBP33 was quantified using a NanoDrop One Microvolume UV-Vis Spectrophotometer (Thermo Fisher Scientific, Waltham, MA, USA), utilizing an absorption coefficient of 0.873 Lg^-1^ cm^-1^ at λ280 nm.

### 2.6. Fluorescence binding assay

N-Phenyl-1-naphthylamine (1-NPN) was used as a fluorescent reporter to assess the ligand binding affinity of LdecOBP33 to various plant volatiles and pesticides, due to the reported success of this probe with insect OBPs in prior studies (Liu et al., 2024; Song et al., 2016). Firstly, to assess the binding affinity of LdecOBP33 to 1-NPN, a saturation binding assay was conducted with a constant protein concentration of 2 µM and a varying concentration of 1-NPN between 0 *µ*M and 250 *µ*M. To assess the binding affinity of various plant volatiles and pesticides to LdecOBP33, competitive fluorescence displacement assays were conducted with LdecOBP33 protein at a constant 2 *µ*M, 1-NPN at 15 *µ*M, and competitor ligands at a concentration varying between 0 *µ*M and 1.8 mM. Each assay was performed in a 96-well flat bottom black plate with a final volume of 200 µl and performed in triplicate. Relative fluorescence intensity was measured with 340nm/35nm excitation filter and a 430 nm/35 nm emission filter, using a multi-mode Tecan Spark® plate reader. GraphPad Prism 8.0 was used to fit the saturation binding curve and to calculate the dissociation constant between LdecOBP33 and 1-NPN, as well as fluorescence inhibition curves to calculate IC_50_ values for competitor ligands. The equation 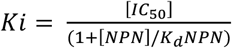 (D’Onofrio et al., 2020) was used to calculate the *K*_*i*_ of the competitor ligands.

### 2.7. LdecOBP33 protein model and ligand docking

The LdecOBP33 protein model was predicted withAlphafold2 using ColabFold v1.5.5: with MMseqs2 (Jumper et al., 2021; Mirdita et al., 2022; Zimmermann et al., 2018). Default settings were used except for relaxations set to 1, template set to pdb100, number of recycles set at 12, the mxa_msa set to 32:64, and with num_seeds set to 2. The full-length minus its signal peptide was used for the AlphaFold2 model. The LdecOBP33 model was superposed with protein-ligand complex crystal structure homologues, that had been identified with HHpred server search; PDB IDs 3R72, 8BXW,100H, to identify the presumptive ligand binding pocket (Kruse et al., 2003; Liggri et al., 2023; Zimmermann et al., 2018). Based on structural alignment with AgamOBP5 (PDB ID 8BXW) co-crystal structure with carvacrol, the C-terminal loop of the LdecOBP33 AlphaFold2 model was remodeled from Ile 131 to Pro 142 with MODELLER and then minimized with Amber ff14SB implemented in Chimera v1.15, before molecular docking was performed (Kruse et al., 2003; Maier et al., 2015; Pettersen et al., 2004; Webb and Sali, 2016). Molecular docking was performed using DockingPie 1.2 plugin installed in PyMOL 3.0.4 with exhaustiveness set to 20, possible poses increased to 3, and grid box was set 20 Å cubed about the presumptive ligand binding pocket, and the scoring function Vinardo with the Smina used (Koes et al., 2013; Rosignoli and Paiardini, 2022). Analysis of the LdecOBP33 and docking was done with Chimera, ChimeraX, and PyMOL, resulting figures were rendered with ChimeraX v1.8 (Goddard et al., 2018; Koes et al., 2013; Meng et al., 2023; Moural et al., 2024; Pettersen et al., 2004; Pettersen et al., 2021).

### 2.8. Bioassay with imidacloprid

Followed by feeding RNAi, control or *LdecOBP33* knockdown CPB male beetles were placed individually into fresh petri dishes and anesthetized on ice for 10 minutes. Afterwards, 0.5 µl of an imidacloprid (Sigma Aldrich, St. Louis, MO, USA) solution with various concentrations in acetone was topically applied to each antenna of CPB using a Hamilton 25 *µ*l model #702 syringe (Hamilton, Reno, NV, USA). Care was taken to ensure that the pesticide solution was applied exclusively to the antennae. A dose of 0.45 *µ*g/µl of imidacloprid solution in acetone was chosen based on our preliminary tests, which determined an LD50 of 0.45 *µ*g/µL for a 0.5 µL application. After the treatment, all CPBs were placed in a fresh petri dish with fresh potato leaves and kept in a rearing room maintained at 25 ± 1°C, 60% ± 5%, and a 16:8 photoperiod regime. CPB mortality was assessed at 0, 3, 6, and 12 hours, and then every 12 hours thereafter for a total of five days post-treatment. The determination of recovery in an individual CPB to pesticide treatment was adapted from prior studies (Mota-Sanchez et al., 2006; Zhao et al., 2000) using three criteria. First, the hind leg was pinched and beetles that failed to respond were considered dead. Second, beetles were placed prone and given two minutes to flip upright; those unable to do so were classified as dead. Finally, beetles were placed upside down on a paintbrush holder and observed; those unable to walk full body length or that fell off were considered dead. Six biological replicates were conducted, each consisting of 8-10 beetles No mortality was observed in beetles treated with acetone alone.

### 2.9. Behavior assay

The ability of CPB adults to locate a host plant was observed in behavioral assays. CPB adults were starved for 24 hours following the dsRNA feeding. Beetles were allotted one hour to acclimate to the room within fresh petri dishes prior to testing. During this time, a 30.50 × 40.65 centimeters aluminum foil sheet was placed within a wind tunnel as the experimental arena, with a constant air flow of 1.5 m/s at 24 ± 2°C and approximately 50% humidity. A potato leaf from a CPB larvae-infested plant served as the odor stimulus and was placed at the upwind edge of the arena, along with a water-saturated cotton ball to maintain constant humidity (which can affect volatile release) (Gouinguené and Turlings, 2002; Schutz et al., 1997b). The leaves were used for 30 minutes before being replaced with fresh infested leaves. The temperature was 22 ± 3 °C, with 50% relative humidity. For the behavior assay, an individual CPB was gently placed at 35 centimeters opposite to the potato leaf and faced upwind. Beetles that walked out of the arena were gently placed back at the starting point. The time taken and ability of each beetle to contact the leaf within 5 minutes was recorded. After the conclusion of each assay the CPB was returned to a fresh petri dish, and the tin foil arena was wiped down with a 70% ethanol solution. Two tin foil arenas were used in total for these bioassays and were used in alternation between individual trials, to ensure any potential semiochemicals host finding ability. Likewise, control and experimentally treated beetles were alternated in bioassays to account for temporal variation. All CPB used were only assayed once and a total of 64 male and 46 female CPB were assayed from both the control and experimental treatment groups.

### 2.10. Electroantennography (EAG) recordings

EAG was used to test the antennal response of control or *LdecOBP33* knockdown CPB to seven potato green leaf volatiles (GLVs), (E)-2-hexenal, (Z)-3-hexen-1-ol, (Z)-3-hexenyl butyrate, 2-phenylethanol, methyl salicylate, linalool, and nonanal (purchased from Sigma Aldrich, St. Louis, MO, USA) that were selected based on previous literature (Dickens, 2000; Visser, 1979; Visser et al., 1979). Samples were prepared to a mass of 1 *µ*g in n-hexane (VWR, Radnor, PA, USA) within a glass scintillation vial, which was subsequently sealed with parafilm and stored at 4°C. Before each bioassay, sample mixtures were brought to room temperature. Odor cartridges were prepared by pipetting 5 µl of each sample mixture on a 10 cm folded strip of filter paper (Whatman No. 1). The filter paper was placed in a disposable glass Pasteur pipette (228 mm long, 7.5 mm wide) after 2 minutes to allow the n-hexane solvent to evaporate, forming an odor cartridge. Antennae of CPB were excised at the base and the pedicel and second to last anterior flagellomere were removed under a dissection microscope. Immediately, the antennae were mounted onto a Quadro-probe (Syntech, Virgina Beach, VA) with electroconductive gel (Physio Control Electrode Gel, Grainger, USA), with the base of the antennae positioned on the reference electrode and the tip on the recording electrode. The Quadro-probe antennal preparation was placed directly next to a dually purified and humidified constant air flow stream maintained at 20 mL/sec in a Faraday cage. Recordings were performed using the Syntech EAG software and began once antennal responses maintained a steady baseline response for at least ten seconds. The antenna was exposed to hexane between each stimulus to assess the diminishing response to stimuli as the antenna aged. Antennae from control and knockdown CPB were alternated between bioassays, to account for temporal variation. The maximum amplitude (mV) that was evoked by an antenna preparation, in response to a test stimulus, was considered as the EAG response. To account for solvent and/or mechanosensory artifacts influencing the EAG response of each antennal preparation, the average of the EAG response to control stimuli was subtracted from the EAG response to the test stimuli (Gabel et al., 1992; Light et al., 1988; Raguso and Light, 1998; Reed et al., 1987). The adjusted average response of each antennal preparation (N = 14) represented the average antennal response to test stimuli in both control and *LdecOBP33* knockdown beetles.

### 2.11. Statistical analysis

Differences in the gene expression across various developmental stages, tissues, and strains were calculated using a one-way ANOVA followed by a Tukey’s HSD test. The differences in the ability to locate their host, detect individual potato plant volatile chemical constituents, and to resist pesticide treatment between the control and *LdecOBP33* knockdown beetles were analyzed using a Student T-Test. To compare the mortality of control and *LdecOBP33* knockdown beetles’ response to 0.5 *u*L of 0.45 *µ*g/µl imidacloprid over time, survival analysis was performed using a log-rank and Wilcoxon test. All statistical analyses were conducted using JMP v 17.0 (SAS Institute, Cary, NC, USA).

## 3. Results

### 3.1. Expression patterns of LdecOBP33 and bioinformatic analyses

Developmental expression showed that *LdecOBP33* was expressed across all development stages, with predominant expression in adult and the highest expression was observed in males (Fig. 1A). The spatial expression pattern of *LdecOBP33* indicated antennal-specific expression, with particularly high levels in male antenna (Fig. 1B).

**Fig. 1.**
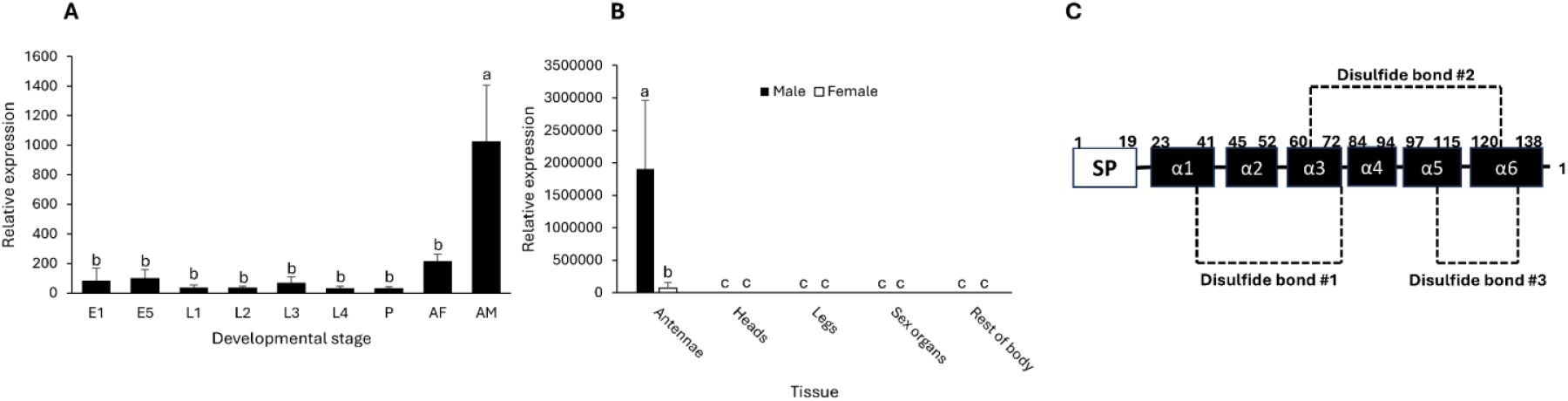
The gene expression patterns, and secondary and quaternary structures of *LdecOBP33*. CPB from each developmental stage (3-50 per, dependent on the state) and specific tissue from one week old adult male and female CPB were used to generate expression pattern analyses. (A) Developmental expressional pattern of *LdecOBP33*. E1, one-day old egg; E5, five-day old egg; L1-L4, first-fourth instar larvae; P, pupae; AF, adult female; AM, adult male (B) Spatial expression pattern of *LdecOBP33*. E1, one-day old egg; E5, five-day old egg; L1-L4, first-fourth instar larvae; P, pupae; AF, adult female; AM, adult male. (C) Schematic diagram of the secondary structure of LdecOBP33. SP, signal peptide; α1-α6, alpha helices 1-6; dashed lines (−-) indicate disulfide bond bridge. Standard error is represented by the error bars; Different alphabetical letters indicate a significant difference of relative *LdecOBP33* gene expression among developmental stages and tissues at p <0.05 (one way ANOVA followed by Tukey’s Test).

LdecOBP33 shared a sequence similarity of 99.53% with an uncharacterized *Leptinotarsa decemlineata* protein (NCBI accession number: XM_023165485). Additionally, the predicted secondary and tertiary structure of LdecOBP33 featured the hallmark characteristics of a classic-class OBP, with six highly conserved cysteines residues and three disulfide bonds (Fig. 1C). The molecular weight of LdecOBP33 was estimated to be at 15.259 kDa, with an isoelectric point of 8.18. A signal peptide between amino acid residues 1-19 was detected as well, which was removed prior to protein expression.

To investigate the evolutionary relationships of LdecOBP33 to other Coleopteran OBPs, we performed a phylogenetic analysis. Following the analysis, it was found that LdecOBP33 resided in one of the two classic OBP clades, alongside two other OBPs with broad binding affinity MaltOBP10 (Li et al., 2020) and MaltOBP13 (Li et al., 2017) (Fig. S1). Specifically, LdecOBP33 clustered together with other insect classic OBPs that also reside in an additional subfamily, termed antennal binding protein II (ABPII) (Andersson et al., 2019; Coates et al., 2023; Dippel et al., 2014).

### 3.2. LdecOBP33 showed a high binding affinity with various plant volatiles and pesticides

To assess the binding affinity of LdecOBP33 to potential ligands, we performed competitive fluorescence displacement assays with LdecOBP33 and various potato plant volatiles and pesticides. Firstly, with highly purified LdecOBP33 protein, we determined the affinity of the fluorescent reporter 1-NPN through a non-linear regression one-site saturation binding curve that accounts for ligand depletion (Motulsky and Neubig, 2010), which resulted in a dissociation constant of 13.41 ± 1.72 *µ*M (Fig. 2A). Secondly, the inhibition of the 1-NPN:LdecOBP33 complex in the presence of competitor ligands was calculated using a non-linear inhibition curve fit model to obtain respective IC_50_ (half-maximal inhibitory concentration) values for each competitor ligand. For plant volatiles, we observed displacement of the 1-NPN probe by over 50% with nonanal (IC_50_ = 18.70 ± 3.32 *µ*M), (Z)-3-hexenyl-butyrate (IC_50_ = 51.50 ± 7.91 *µ*M), L-linalool (IC_50_ = 78.62 ± 9.24 *µ*M), (E)-2-hexenal (IC_50_ = 405.60 ± 14.84 *µ*M), methyl salicylate (IC_50_ = 1128 ± 125.30 *µ*M), (Z)-3-hexen-1-ol (IC_50_ = 1176 ± 181.40 *µ*M), and 2-phenylethanol (IC_50_ = 1304 ± 226.50 *µ*M) (Table 1, Fig. 2B&C). For pesticides tested, displacement of the 1-NPN probe by over 50% was observed with tetramethrin (IC_50_ = 40.47 ± 2.40 *µ*M), clothianidin (IC_50_ = 40.49 ± 6.63 *µ*M), chlorpyrifos-methyl (IC_50_ = 272.80 ± 21.64 *µ*M), and imidacloprid (IC_50_ = 317.20 ± 35.74 *µ*M) (Table 1, Fig. 2D), indicating that this OBP has a rather broad binding spectrum. No noticeable displacement of the 1-NPN probe in the presence of the permethrin, cypermethrin, fenvalerate, carbaryl, and myclobutanil were observed (Table 1).

**Table 1.** Binding data of competitor ligands to LdecOBP33. Note: Competitor ligands that did not display any intrinsic binding affinity to LdOBP33 are denoted with a dash (−) in the table

| Category | Ligand name | IC50 (μM) | Ki (μM) | LogP |
| --- | --- | --- | --- | --- |
| Potato plant volatile | Nonanal | 18.70 ± 3.32 | 9.15 | 3.30 |
|  | (Z)-3-hexenyl-butyrate | 51.50 ± 7.91 | 25.21 | 2.70 |
|  | L-Linalool | 78.62 ± 9.24 | 38.48 | 2.70 |
|  | (E)-2-hexenal | 405.60 ± 14.84 | 198.53 | 1.50 |
|  | Methyl salicylate | 1128 ± 125.30 | 552.13 | 2.50 |
|  | (E)-3-hexen-1-ol | 1176 ± 181.40 | 575.62 | 1.30 |
|  | 2-Phenylethanol | 1304 ± 226.50 | 638.28 | 1.36 |
| Plant product | D-Glucose | - | - | -2.60 |
| Neonicotinoid | Clothianidin | 40.49 ± 6.63 | 19.82 | 0.70 |
|  | Imidacloprid | 317.20 ± 35.74 | 155.26 | 0.57 |
| Organophosphate | Chlorpyrifos-methyl | 272.80 ± 21.64 | 133.53 | 4.30 |
| Carbamate | Carbaryl | - | - | 2.36 |
| Pyrethroid | Tetramethrin | 40.47 ± 2.40 | 19.81 | 4.70 |
|  | Permethrin | - | - | 6.50 |
|  | Cypermethrin | - | - | 6.60 |
|  | Fenvalerate | - | - | 6.20 |

**Fig. 2.**
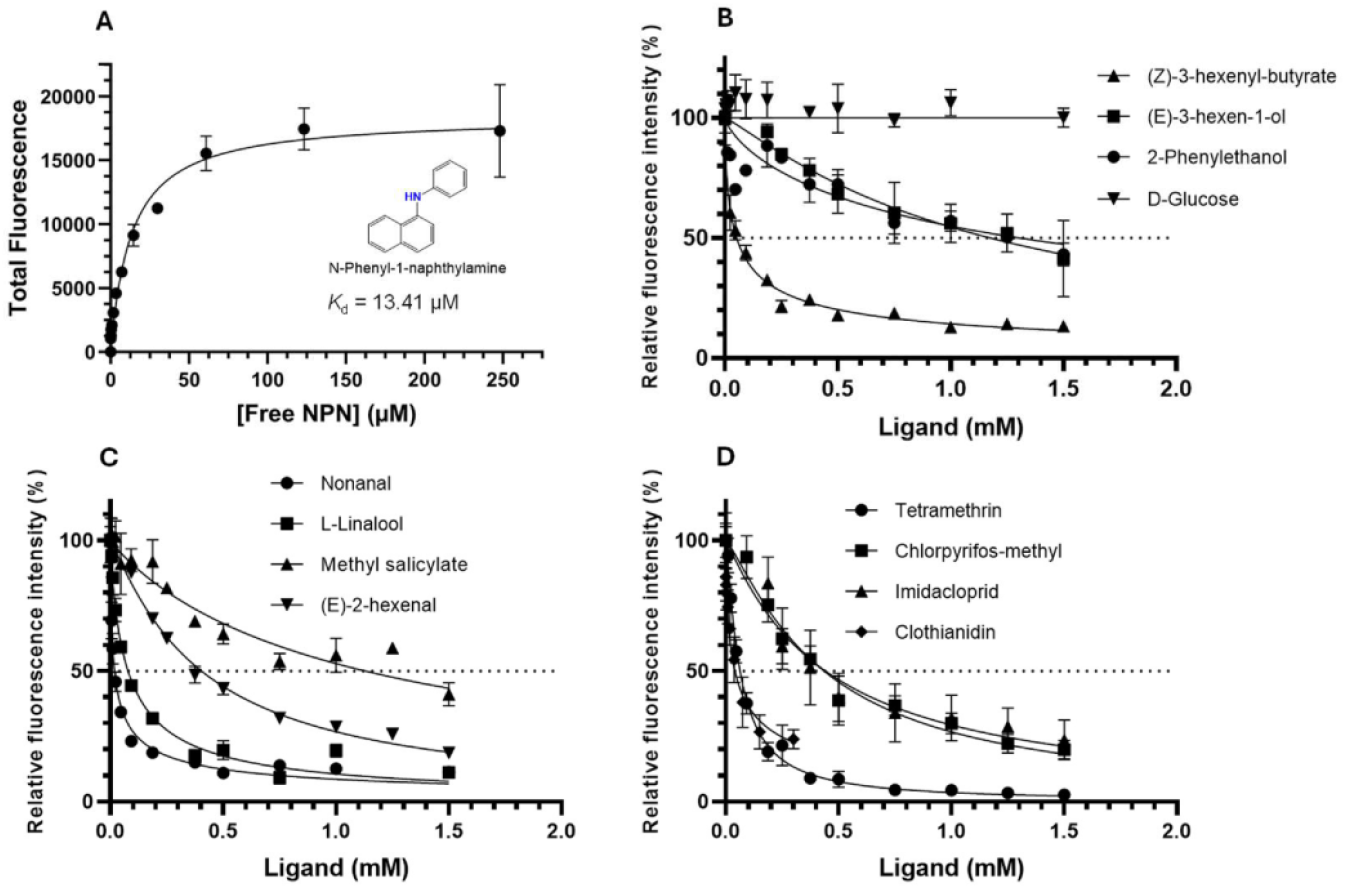
AlphFold2 model of LdecOBP33. (A) The global fold of LdecOBP33 displayed as a ribbon diagram (helices colored blue and coils colored tan) overlayed with transparent surface showing ligand binding pocket. The conserved are shown and labeled with their respective cysteine residues. (B) Highest scoring pose of imidacloprid (colored by heteroatom, with carbons colored cyan), all sidechains within 4 Å are displayed and labeled. (C) Highest scoring pose of imidacloprid (colored by heteroatom, with carbons colored green), all sidechains within 4 Å are displayed and labeled.

Using the respective IC_50_ value obtained for each competitor ligand, the inhibitory constant (*K*i) was calculated with the Cheng and Prusoff equation (Cheng and Prusoff, 1973). The calculated Ki values for the plant volatiles were 9.15 µM for nonanal, 25.21 *µ*M for (Z)-3-hexenyl-butyrate, 38.48 *µ*M for L-linalool, 198.53 *µ*M for (E)-2-hexenal, 552.13 *µ*M for methyl salicylate, 575.62 *µ*M for (Z)-3-hexen-1-ol, and 638.28 *µ*M for 2-phenylethanol. The calculated Ki values for pesticides were 19.81 *µ*M for tetramethrin, 19.82 *µ*M for clothianidin, 133.53 *µ*M for chlorpyrifos-methyl, and 155.26 *µ*M for imidacloprid (Table 1).

### 3.3. Structure modeling and molecular docking with nonanal and imidacloprid

The AlphaFold2 model of LdecOBP33 revealed a compact bundle of six α-helices surrounding a hydrophobic ligand binding pocket (Fig. 3A). The fold is stabilized by three conserved disulfide bonds, formed by Cys37-Cys68, Cys64-121, and Cys110-Cys130. Structural alignment with homologues identified the ligand binding pocket. As depicted in Fig. 3A, the ligand binding pocket opens near the N-terminus and extends as a channel running toward the C-terminus. While many residues lining the pocket are hydrophobic, some polar and charged residues are also present and are expected to play key roles in hydrogen bonding with ligands. Molecular docking with nonanal and imidacloprid revealed two key residues potentially involved in hydrogen bond formation. In the highest scoring pose (−5.9 kcal/mol) of imidacloprid, a hydrogen bond (2.15 Å) between Asn138 HD2 and the O2 atom of imidacloprid was observed (Fig. 3B). Additionally, in the top ranked pose of nonanal, the carbonyl oxygen formed a hydrogen bond (3.05 Å) with the HD21 hydrogen of Asn94 (Fig. 3C).

**Fig. 3.**
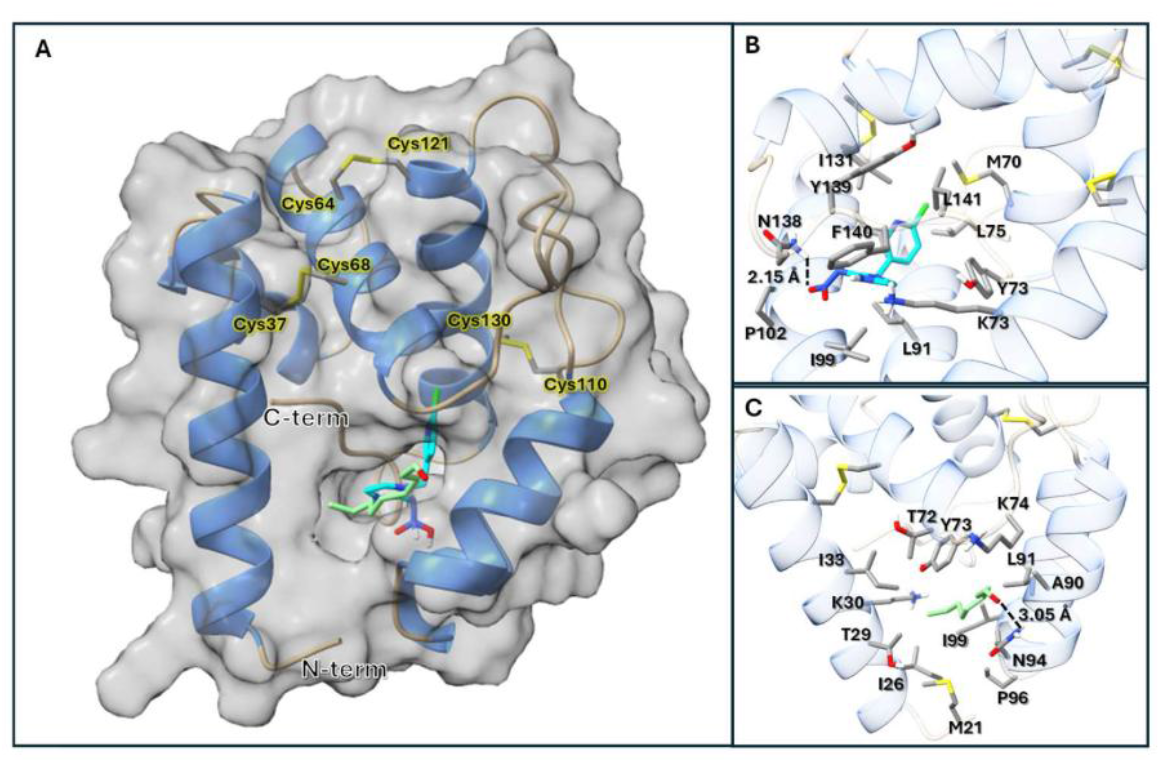
Competitive binding curves of plant compounds and pesticides to purified recombinant LdecOBP33 protein. (A) Affinity of purified recombinant LdecOBP33 protein to the fluorescent reporter 1-NPN. Kd, dissociation constant. (B,C,D) Affinity of potato plant compounds and pesticides to purified recombinant LdecOBP33 protein. The dashed line is indicative of 50% displacement of the 1-NPN probe. Curves are resultant of 2-3 independent replicates.

### 3.4. LdecOBP33 contributed to imidacloprid resistance in CPB

Due to the robustness of CPB to develop resistance to pesticides, along with the high binding affinity of LdecOBP33 towards various insecticides, we investigated whether *LdecOBP33* exhibited differential expression between the susceptible and imidacloprid-resistant strains of CPB. Interestingly, we found that *LdecOBP33* expression was 2.5-fold higher in the resistant strain compared to the susceptible one (Fig. 4A). To determine whether *LdecOBP33* contributed to imidacloprid resistance in CPB, we first silenced *LdecOBP33* expression in the imidacloprid-resistant CPB males by feeding RNAi. After 5 days, *LdecOBP33* showed a 77% reduction in transcriptional expression level in ds*LecOBP33* feeding beetles compared to *dsGFP* feeding beetles, confirming the successful silencing of *LdecOBP33* (Fig. 4B). Next, we conducted an antenna contact bioassay by applying imidacloprid on both antennae of CPB males (Fig. 4C) and monitored mortality over five days. During the first 36 hours, there was no significant difference in mortality between the *LdecOBP33* silencing group and the ds*GFP* control group (Fig. 4D). However, after 36 hours, the ds*LdecOBP33*-KD beetles showed a significant higher mortality after exposure to imidacloprid in ds*LdecOBP33*-KD beetles (81%) compared to the control beetles (55%) (Fig. 4D), indicating a general decrease in survival. By 120 hours, the average mortality (N = 52, 6 replicates) in ds*LdecOBP33* treated CPB was 32% while the mortality of control was 12%, increasing by 19% (p-value < 0.001) (Fig. 4D).

**Fig. 4.**
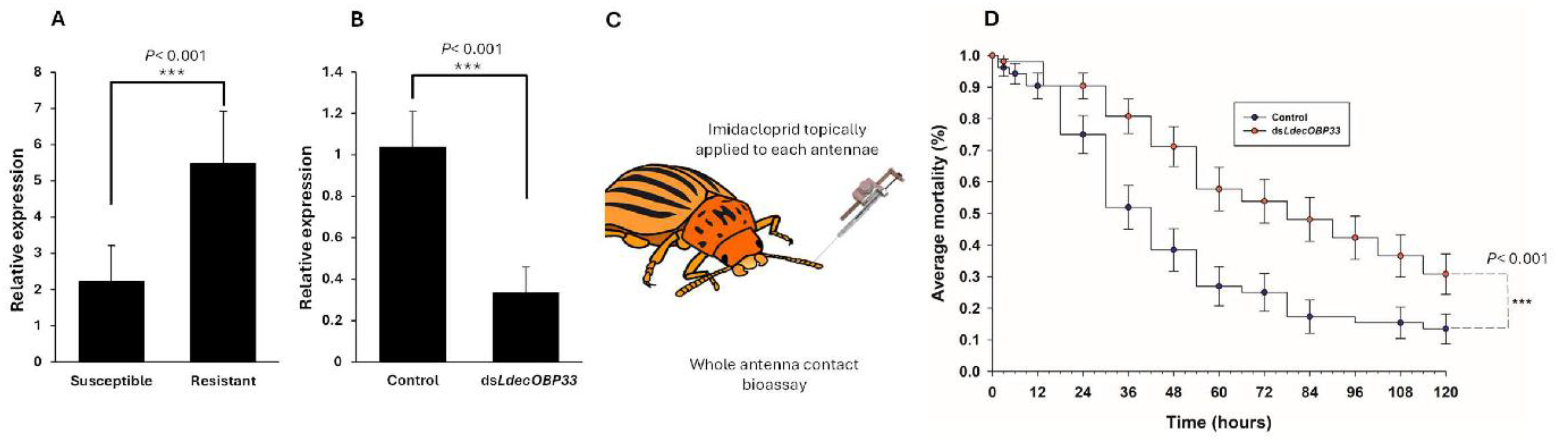
Influence of *LdecOBP33* towards pesticide resistance in adult male CPB. (A) Differential expression of *LdecOBP33* in a neonicotinoid-susceptible and resistant strain of CPB. (B) Relative expression of *LdecOBP33* in CPB fed ds*GFP* (control) and ds*LdecOBP33*. A p-value < 0.001 (Student *t*-test) is denoted as “***” and represents a difference in the expression of *LdecOBP33* between the susceptible and resistant strain of CPB (A) or between the control and knockdown treatments (B). (C) Cartoon for the whole antenna contact bioassay. CPB were anesthetized on ice for 10 minutes and were then directly administered 0.50 *µ*g/µl of an imidacloprid solution topically to each antenna. (D) Influence of dsLdecOBP33 on adult male CPB survival across 120 hours (five days). A p-value < 0.001 (Log-Rank & Wilcoxon Test) is denoted as “***” and represents a difference in the survival of adult male CPB who survived a treatment of 0.45 *µ*g/*µ*l pesticide treatment (imidacloprid diluted in acetone) topically applied to both antennae, between the control and knockdown groups.

### 3.5. Roles of LdecOBP33 in host location

We compared the response of *LdecOBP33-*KD beetles and control beetles to potato leaves infested with CPB larva in the wind tunnel to investigate the behavioral role of *LdecOBP33* in host location (Fig. 5A). We recorded two key aspects of the ability of host location: 1). Whether beetles located their host odor source (an excised leaf) within the 5-minute test period and 2) the time that was taken to contact the leaf. After *LdecOBP33* was silenced, the host location ability of adult male CPB declined. Specifically, significantly fewer male beetles in the *LdecOBP33* silenced group successfully located the potato leaf within the 5-minute test period, compared to male beetles in the control group (Fig. 5B). However, no difference was observed in female beetles (Fig. 5C). Among the beetles that reached the potato leaves within 5 minutes, the *LdecOBP33* silenced male beetles took significantly longer to arrive than the control beetles (*p*-value < 0.001, N=64) (Fig. 5D). On average, the *LdecOBP33* silenced male beetles contacted the leaf in 205 seconds (95% CI: 176 to 234 seconds), whereas control beetles took 106 seconds (95% CI: 81 to 132 seconds) (Fig. 5D). We did not observe significant difference in the time of female CPBs took to reach the leaves between *LdecOBP33-*silenced and the control female beetles (Fig. 5E).

**Fig. 5.**
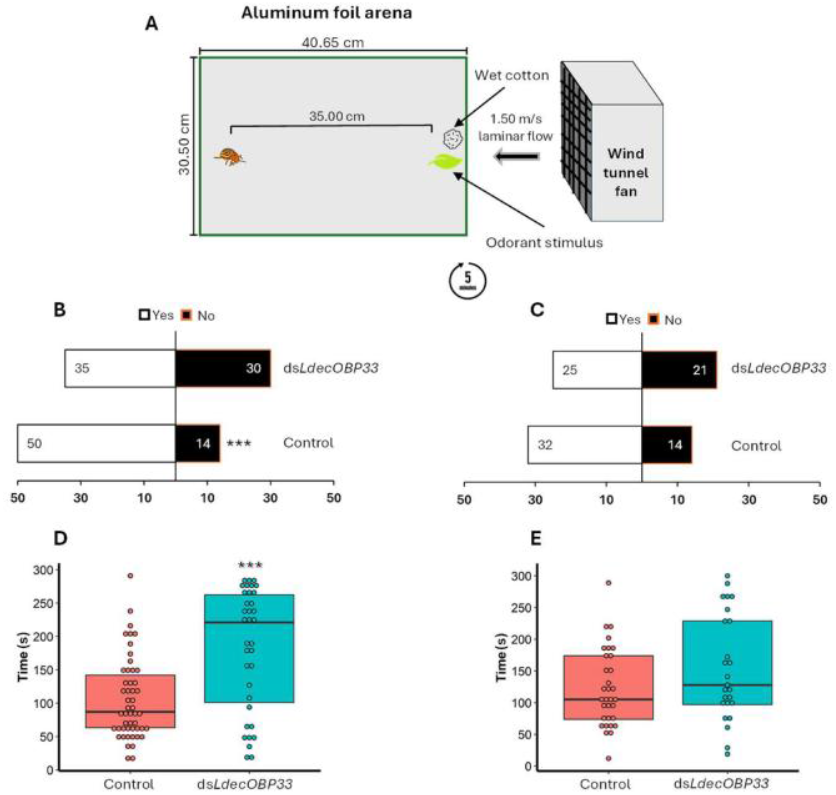
Influence of *LdecOBP33* on host locating capabilities in adult CPB. (A) Depiction of the setup used to perform behavioral assays designed to assess the ability of *LdecOBP33* towards host plant location. CPB were placed on a tin-foil arena within a wind tunnel and given five minutes to locate potato leaf tissue. Each CPB was given 5 minutes for the trial. (B,C) Difference in the ability of adult male (B) and female (C) CPB to locate potato leaves within the five-minute test period, following gene-silencing of *LdecOBP33*. (D,E) Difference in the time of adult male (D) and female (E) CPB to successfully locate the potato leaves within the five-minute test period, following gene-silencing of *LdecOBP33*. A p-value < 0.01 is denoted as “**” and a p-value < 0.001 is denoted as “***” (ANOVA & Student T-test), in which represents a difference in the time and/or ability of CPB to locate their host between the control and knockdown treatment groups. Individual circles on figure B indicate time an individual CPB found the potato leaf.

### 3.6. Impacts of LdecOBP33 silencing on detection of potato plant volatiles

To supplement the behavioral assay, we performed EAG to determine if *LdecOBP33* silencing influenced the antennal response of adult male CPB to previously identified antennal active volatiles from potato plant. The normalized average antennal EAG response to (Z)-3-hexenyl butyrate, (Z)-3-hexen-1-ol, 2-phenylethanol, and methyl salicylate decreased following gene silencing of *LdecOBP33* in CPB males (Fig. 5). There was no difference in the antennal response of adult male CPB to (E)-2-hexenal, nonanal, or linalool detected (Fig. 6). In addition, there was no difference to all potato volatiles tested in CPB females (Fig. S2).

**Fig. 6.**
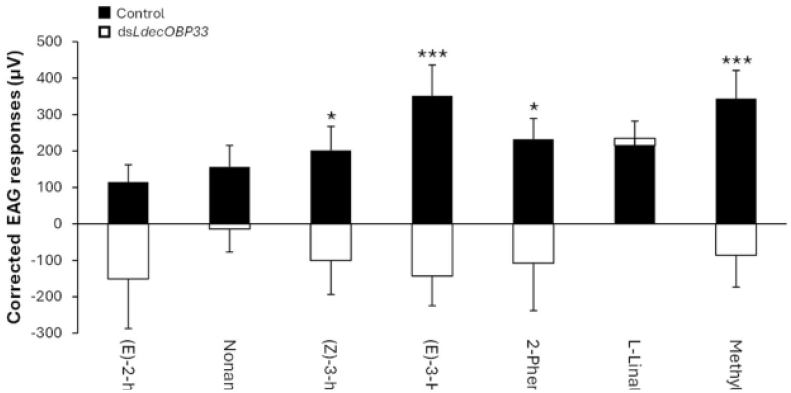
Effect of ds*LdecOBP33* on the average antennal response of adult male CPB to major chemical constituents of the potato plant volatile blend. Antenna were excised and mounted onto a Quadro-probe, wherein they were exposed to 5 *µ*g/*µ*l stimulus loads of potato plant volatiles diluted in hexane. EAG responses of CPB antennae to volatiles were corrected by subtracting the EAG response of the standard (hexane individually) from the EAG response to the potato plant volatiles. A p-value < 0.05 is denoted as “*” and a p-value < 0.001 (Student *t*-test) is denoted as “***”, in which represents a difference in the average antennal response of adult male CPB to a volatile between the control and knockdown groups.

## 4. Discussion

We conducted a comprehensive functional analysis of an antenna-specific OBP in CPB, revealing its role in linking olfaction and xenobiotic adaptation in this species. *LdecOBP33* is most highly expressed in the antennae of male CPB and belongs to the subfamily of antennal binding protein II (ABPII), the classic OBPs. Recombinant LdecOBP33 exhibited a broad binding spectrum, similar to many other insect classic OBPs (Ma et al., 2023; Zhang et al., 2023; Zhou et al., 2024). LdecOBP33 exhibited a preference for binding to the most hydrophobic molecules among those that were tested in this study, following a general trend of an increase in affinity to a molecule as the lipophicity (LogP) value increased. Classically, OBPs have been understood to play a key role in transporting exogenous hydrophobic molecules through the aqueous sensillar lymph (Leal, 2013; Pelosi et al., 2018; Sun et al., 2018). However, recent research strongly suggested that OBPs may also function as a scavenger, interacting with excessive molecules in the perireceptor space, either clearing insoluble ligands that persist or modulating the amount that reaches olfactory receptors (Larter et al., 2016; Pelosi et al., 2018; Wang et al., 2020). Our binding assays suggest that LdecOBP33 exhibits broad affinity and tuning of LdecOBP33 towards hydrophobic molecules, such as potato volatiles and pesticides, potentially facilitating their rapid processing in the perireceptor space and protecting the insect nervous system from harmful effects.

Serving as the initial interface between the environment and the insect olfactory system, OBPs solubilize and transport hydrophobic odorant molecules through the sensillar lymph to olfactory receptors (Leal, 2013). Pesticides may penetrate the antennal lymph and be sequestered by OBPs, preventing their interaction with target proteins and mitigating potential neurotoxic effects (Pu and Chung, 2024). Thus, OBPs are hypothesized to play a role in insecticide resistance. Previous studies have shown that certain insect OBPs can bind pesticides, and some OBPs are upregulated in insecticide resistant strains or in response to insecticide exposure (Liu et al., 2024). Moreover, RNAi-mediated silencing of specific OBPs has been associated with increased insect susceptibility to insecticides (Shen et al., 2022; Zhang et al., 2022). In this study, differential gene expression analysis of *LdecOBP33* revealed that expression was 2.5-fold higher in a neonicotinoid resistant strain in comparison to the susceptible strain. RNAi-mediated gene silencing of *LdecOBP33* led to increased imidacloprid susceptibility in resistant beetles, suggesting that *LdecOBP33* contributes to imidacloprid resistance. Similarly, research has shown that chemosensory proteins also contribute to insecticide resistance, suggesting a broader role for insect odorant and chemical-binding proteins in detoxification mechanisms (Ingham et al., 2020; Pelosi et al., 2018; Pu and Chung, 2024).

It is understood that behavioral adaptation of an insect to their host can be influenced by the evolution of their chemosensory system (Bruce et al., 2005; Chapman, 2003; Jacquin-Joly and Merlin, 2004; Jermy and Szentesi, 2003). In comparison to other chemosensory gene families, OBPs are highly divergent in both form and function, exhibiting low sequence similarity even within members of the same species (Abendroth et al., 2023; Rihani et al., 2021). Due to their important role as the primary mediator between the external environment and the insect olfactory system, OBPs may play an important role in adaptation of insects to their host plants. Prior studies have observed that CPB can discriminate their preferred host plant potato, from other solanaceous species by detecting a specific ratio of volatile compounds unique to potato plants (Hitchner et al., 2008; Visser and Ave, 1978; Visser et al., 1979). Disruption of CPB’s ability to properly perceive odor blend ratios of potato can impair their ability to locate their host plant. As demonstrated in prior studies. Upon modification of the potato odor blend, or the intermixing of other plants with potato, positive anemotaxis behavior of CPB towards potato plants is lost (Jermy et al., 1988; Thiery and Visser, 1986, 1987). In this study, we found that after knockdown of *LdecOBP33* adult male CPBs took significantly longer to locate their host plant and had reduced antennal responses to four potato plant volatiles, compared to the control group. One of these volatiles, (Z)-3-hexen-1-ol, is a major component of the potato plant odor blend and elicits strong antennal responses in adult CPB (Dickens, 2000; Visser, 1979; Visser et al., 1979). The other three volatiles, Z-3-hexenyl-butyrate, 2-phenylethanol, and methyl salicylate, are plant volatiles that can be emitted under stress and both attract and elicit antennal responses from CPB (Dickens, 2000; Schutz et al., 1997a). Moreover, methyl salicylate specifically has been used in several synthetic blends that have elicited attractive response from CPB (Dickens, 2002; Hammock et al., 2007). These findings are consistent with previous studies showing that RNAi-mediated gene silencing of insect OBPs influence host location and antennal responses to host odorants (Biessmann et al., 2010; Li and Zhang, 2022; Tan et al., 2023).

Additionally, in this study, we observed that recombinant LdecOBP33 protein exhibited broad binding affinity to many potato plant volatiles. Volatiles that displayed the greatest affinity to LdecOBP33 were nonanal, (Z)-3-hexenyl-butyrate, and L-linalool, all of which were the most hydrophobic (LogP) volatiles assayed. In contrast to our findings in the EAG bioassay, we found that only one of the four potato plant volatiles ((Z)-3-hexenyl-butyrate) that elicited weaker antennal responses, in silenced beetles compared to control beetles, had strong binding affinity to recombinant LdecOBP33 protein. We observed affinity in the other three volatiles, ((E)-3-hexen-1-ol, methyl salicylate, 2-phenylethanol) as they were able to displace the 1-NPN fluorescent probe by over 50%, but only at a much higher concentration of ligand in the millimolar range, as compared to the low micromolar range. A similar instance of this phenomena was also observed in recent studies (Li and Zhang, 2022; Zhao et al., 2021). These suggest there may not necessarily be a positive correlation between the binding affinity of an OBP to an odorant and the ability of an odorant to induce olfactory activity. Nonetheless, the broad affinity of LdecOBP33 to many potato plant volatiles, alongside our findings from the *in vivo* and *in vitro* analyses, suggests that *LdecOBP33* is fundamental in olfactory processing in adult male CPB.

In conclusion, our study is the first to functionally characterize the roles of any OBP in CPB. We found that *LdecOBP33*, a male antenna-specific OPB, plays dual roles in host location and pesticide resistance. Our findings highlight a potential mechanism that is involved in the response of CPB to environmental stressors and challenges, supporting a potential link between insect adaptation to xenobiotics and olfactory processing.

## Acknowledgements

This work was supported by NSF CAREER IOS-2144082, the USDA National Institute of Food and Federal Appropriations under Hatch Project #PEN04770 and Accession #1010058 (to FZ). TM was supported by USDA NIFA postdoctoral fellowship, grant #2020-67034-31780/project accession #1022959 (2020-2022) and USDA NIFA Hatch Project #PEN04897 and Accession #7005652. We are grateful to Drs. Blair Siegfried, Rudolf Schilder, and Christina Rosa (The Pennsylvania State University) for their suggestions and comments on the project.

---

**Fig. S1.**
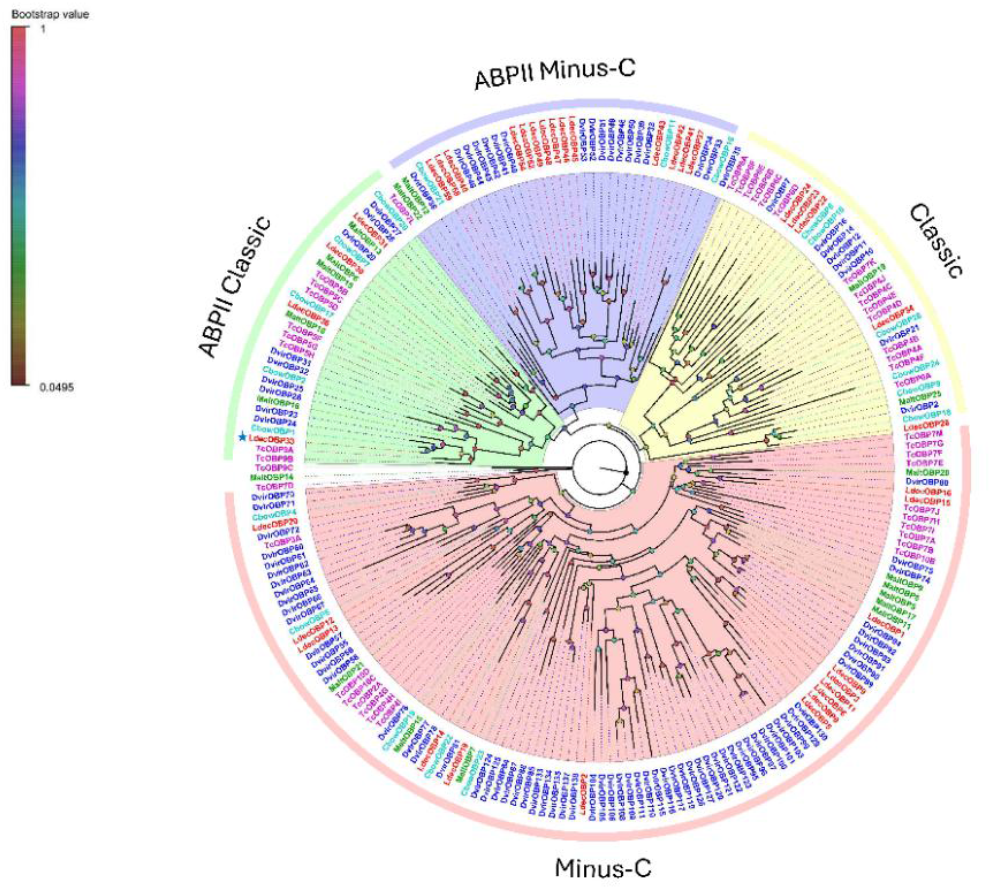
Phylogenetic relationships of Colorado potato beetle (CPB) odorant binding proteins (OBPs) across different coleopteran species. Yellow (classic OBPs), green (ABPII classic OBPs), red (minus-C OBPs), or purple (ABPII minus-C OBPs) coloration indicates the classes of OBPs. Color of label text indicates species (blue, *Diabrotica virgifera virgifera*; purple, *Tribolium castaneum*; light blue, *Collaphellus bowringi*; red, *Leptinotarsa decemlineata*; green, *Monochromatus alternatus*). Star denotes the location of LdecOBP33. The phylogenetic tree was inferred using the maximum likelihood estimation method, with the LG + G + I model, using MEGA11 software. The tree was visualized using Figtree v1.4.4 software.

**Fig. S2.**
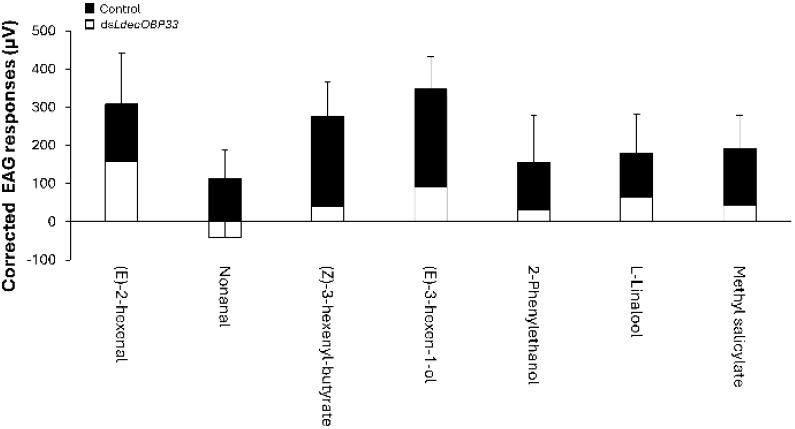
Effect of ds*LdecOBP33* on the average corrected antennal response of female CPB to major chemical constituents of the potato plant volatile blend. Antenna were excised and mounted onto a Quadro-probe, wherein they were exposed to 5 *µ*g/µl stimulus loads of potato plant volatiles diluted in hexane. EAG responses of CPB antennae to volatiles were corrected by subtracting the EAG response of the standard (hexane individually) from the EAG response to the potato plant volatiles.

**Table S1.**
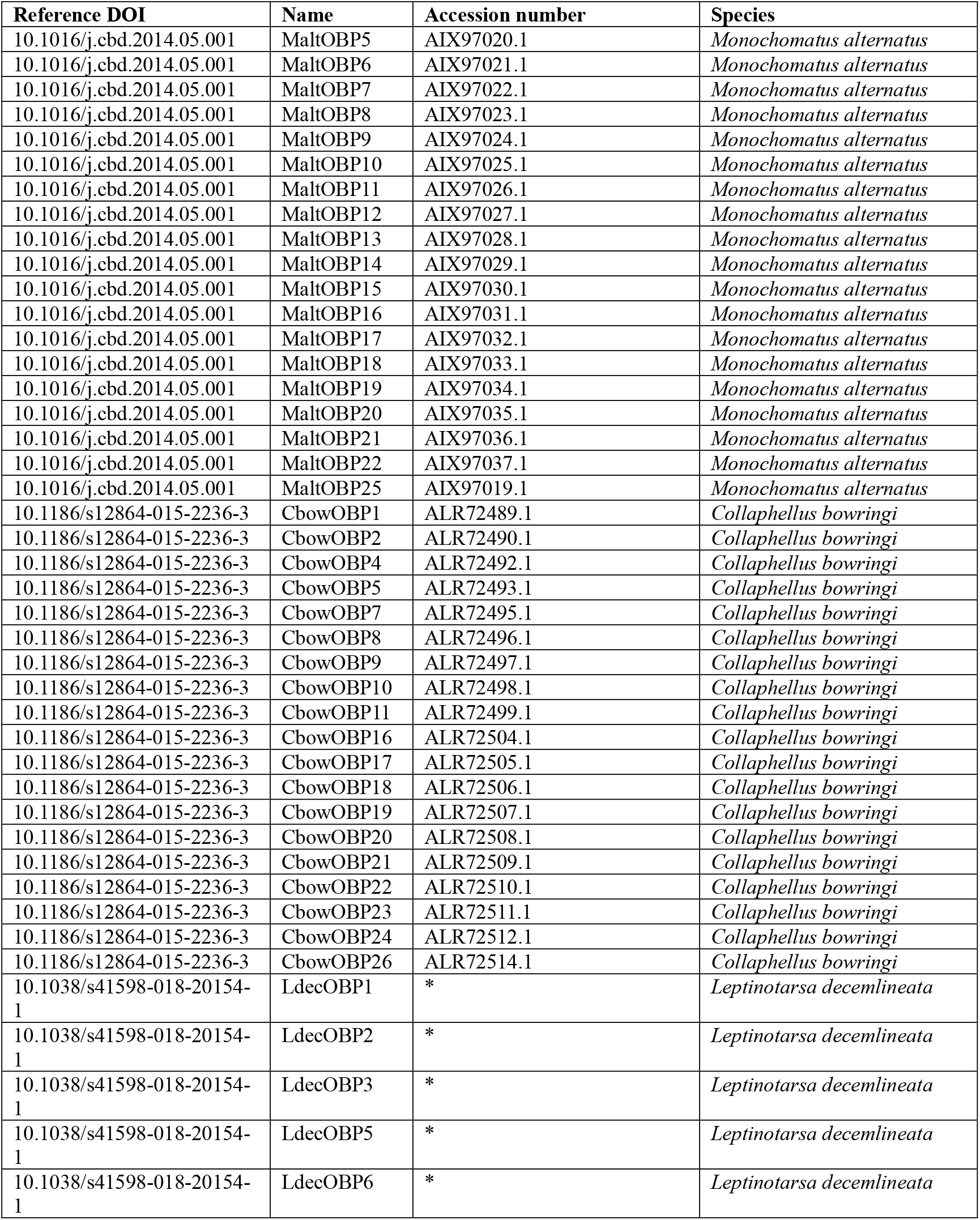
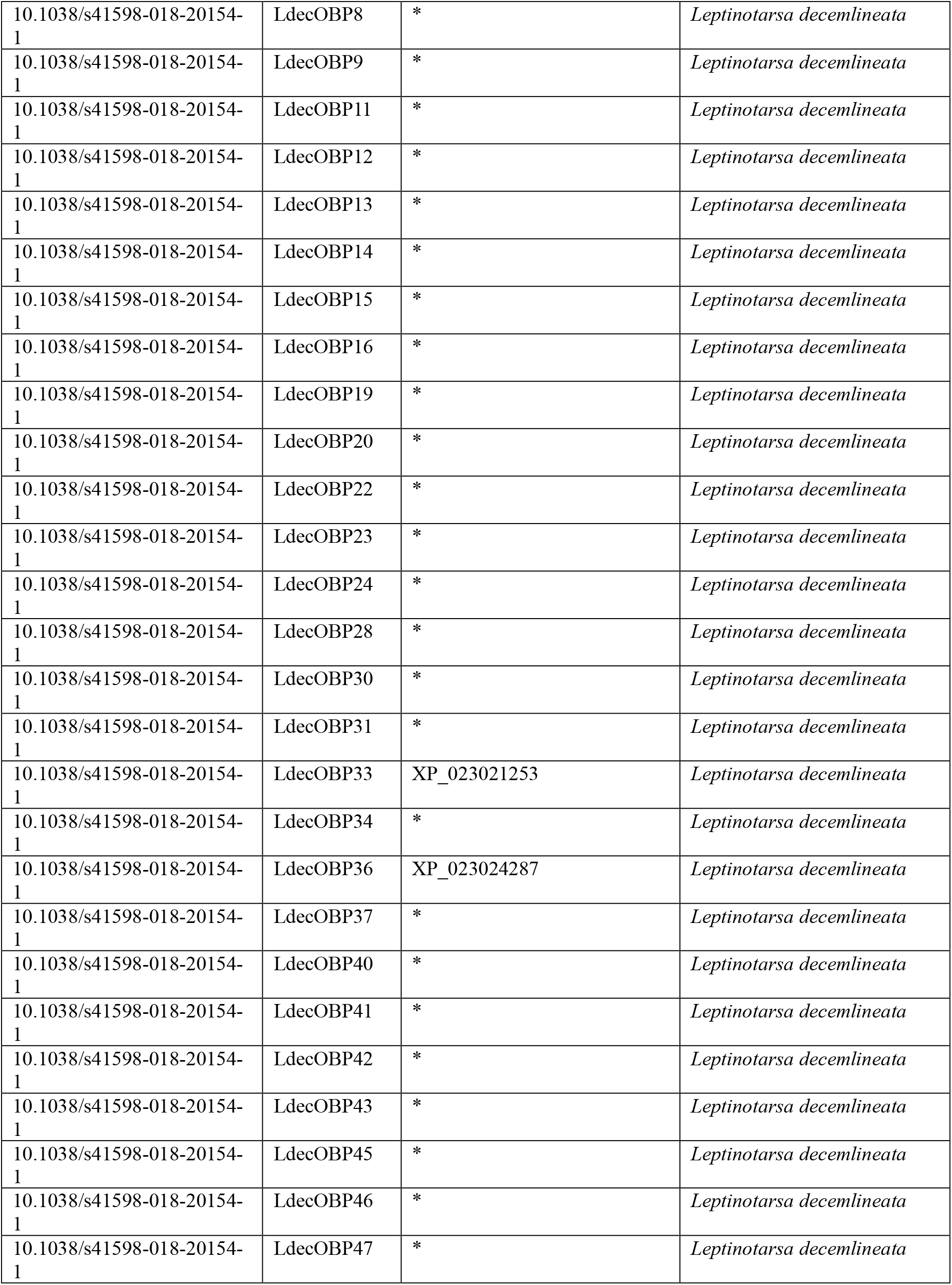
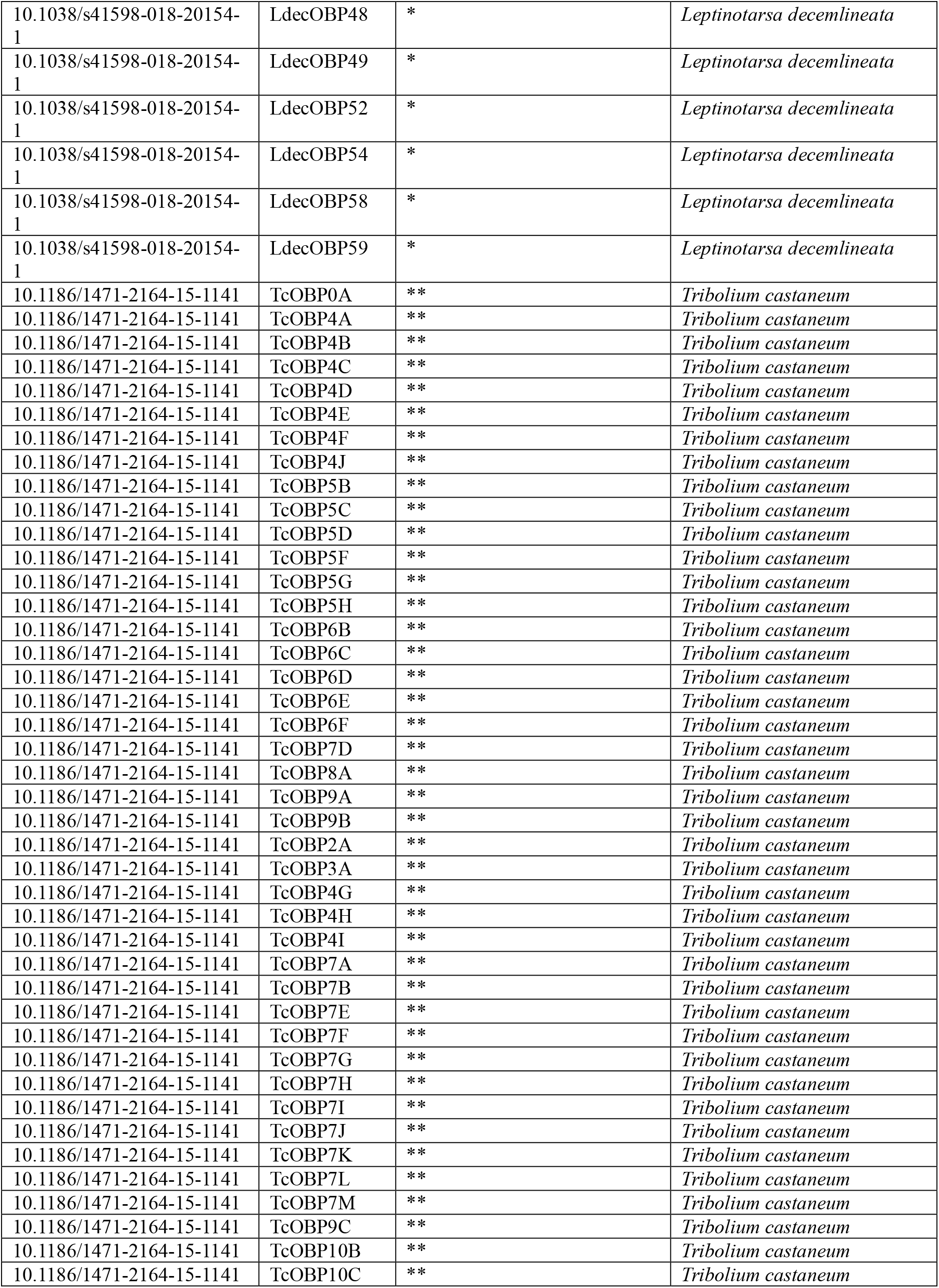
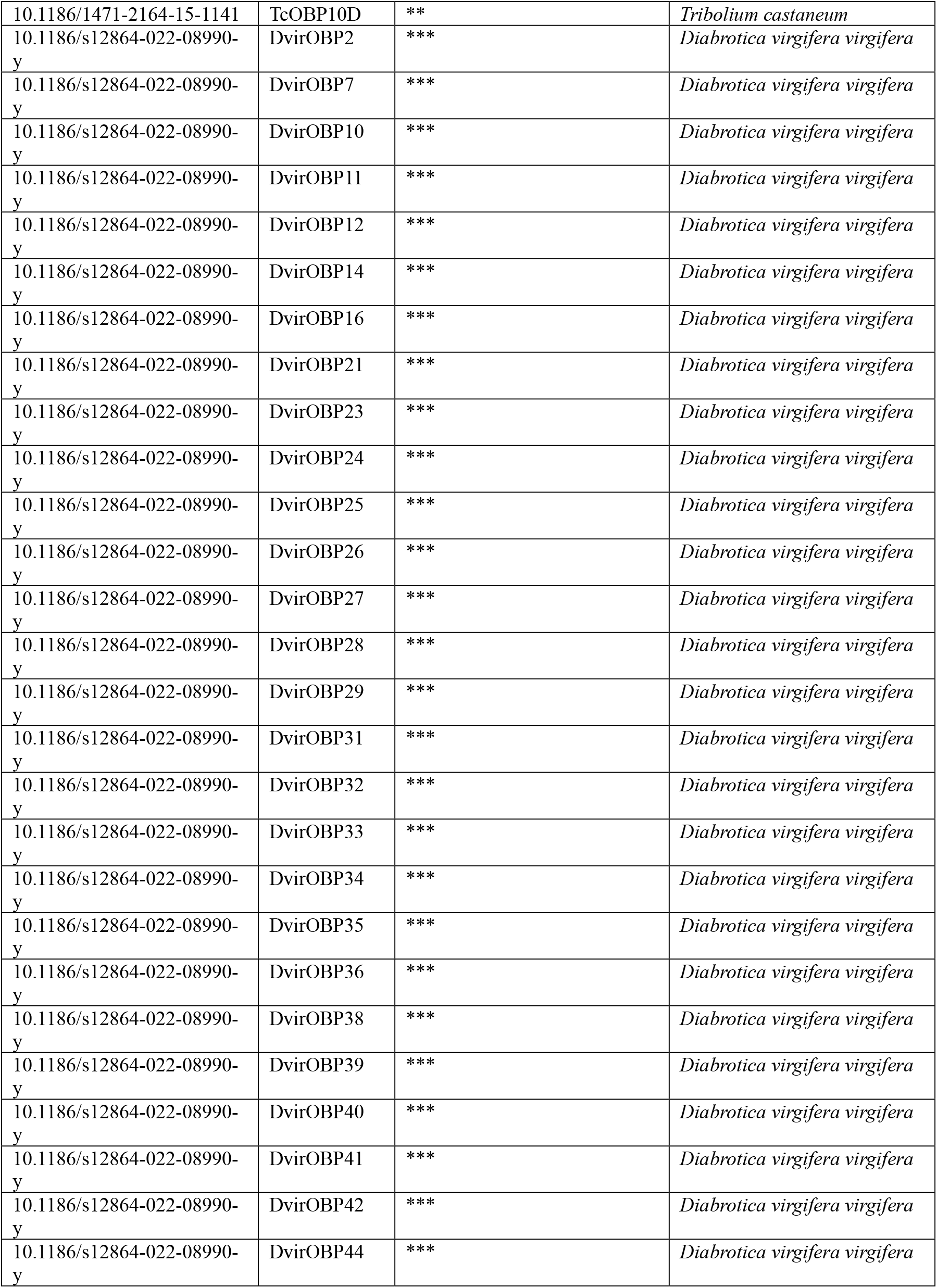
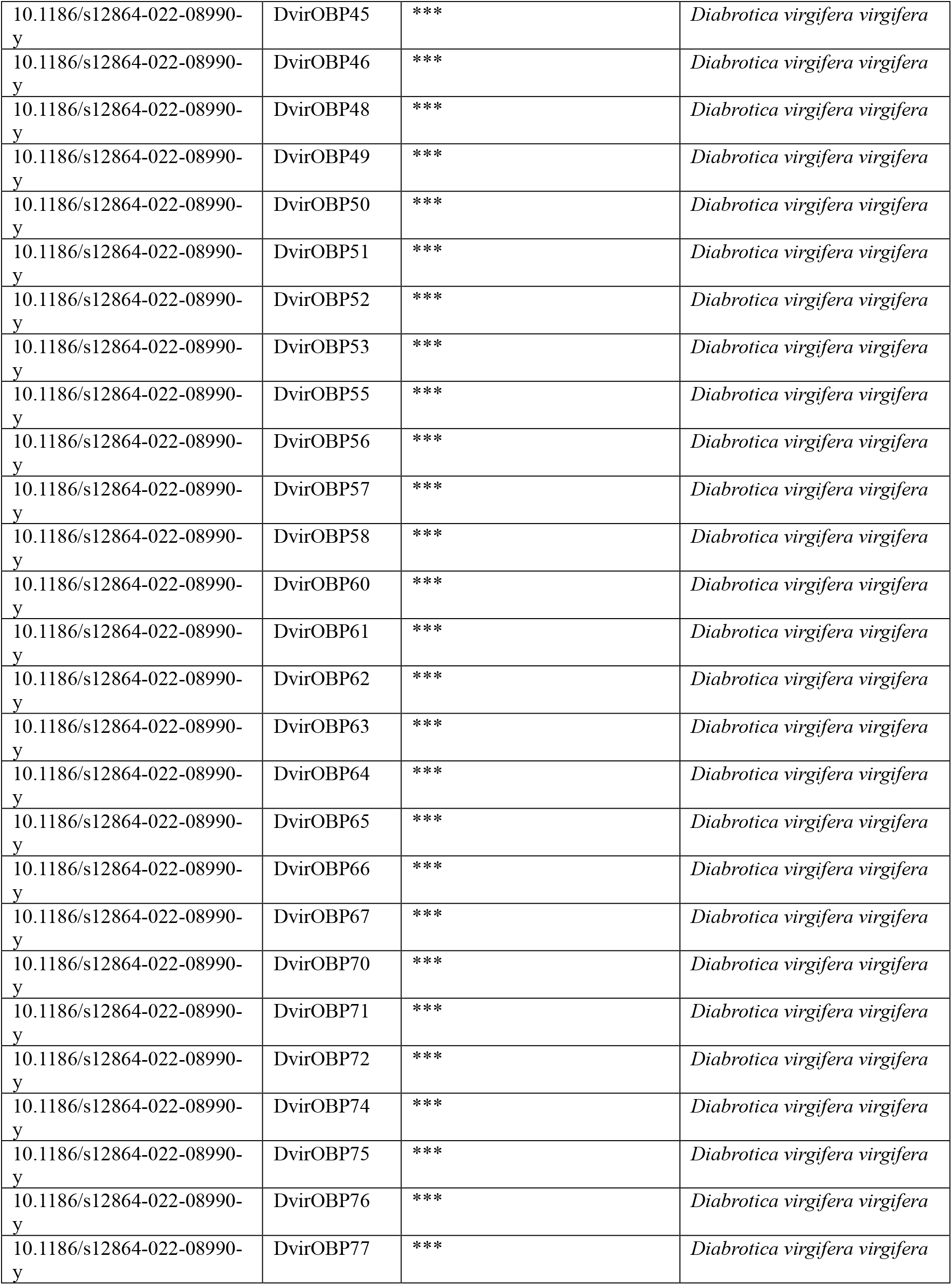
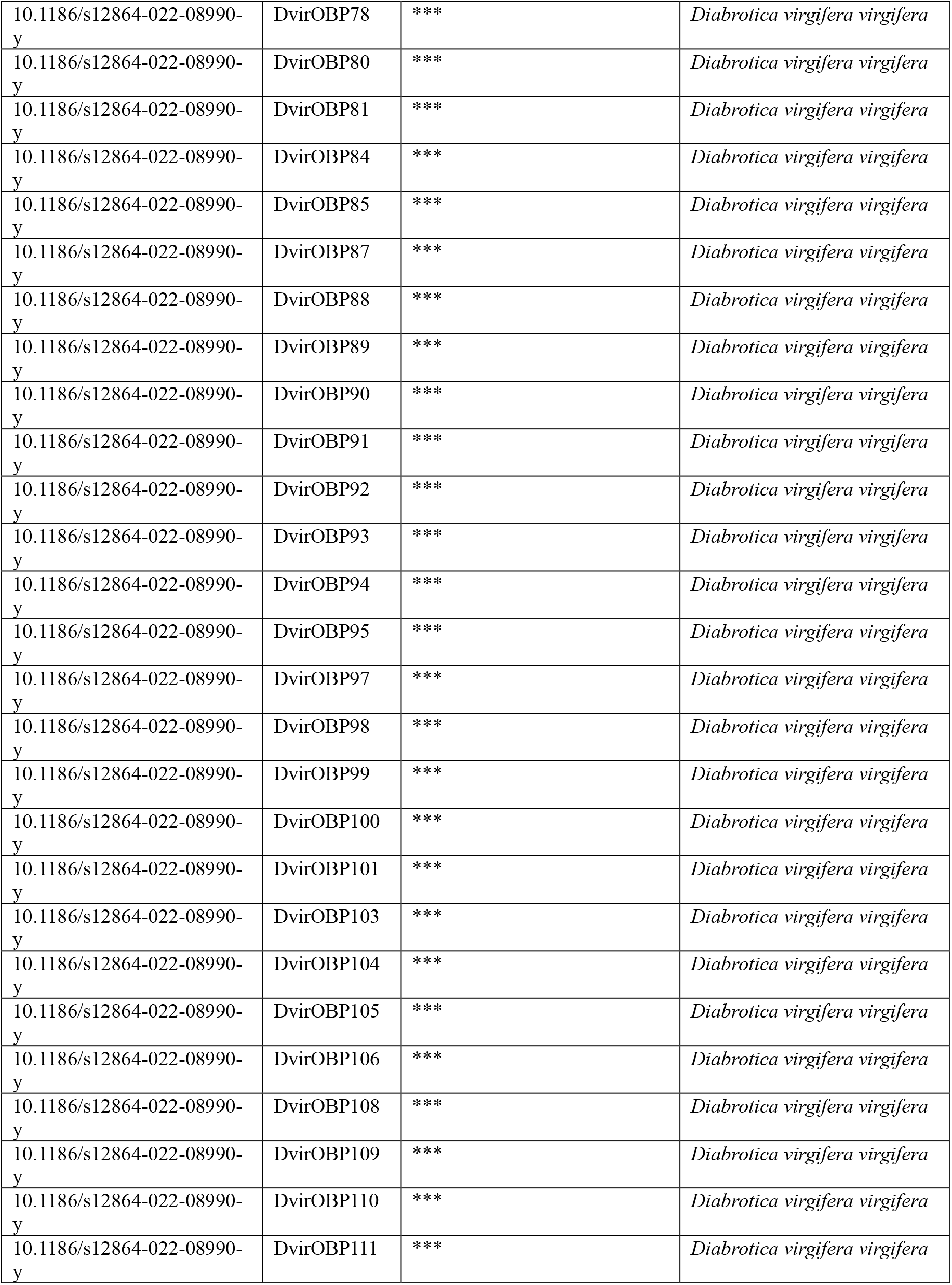
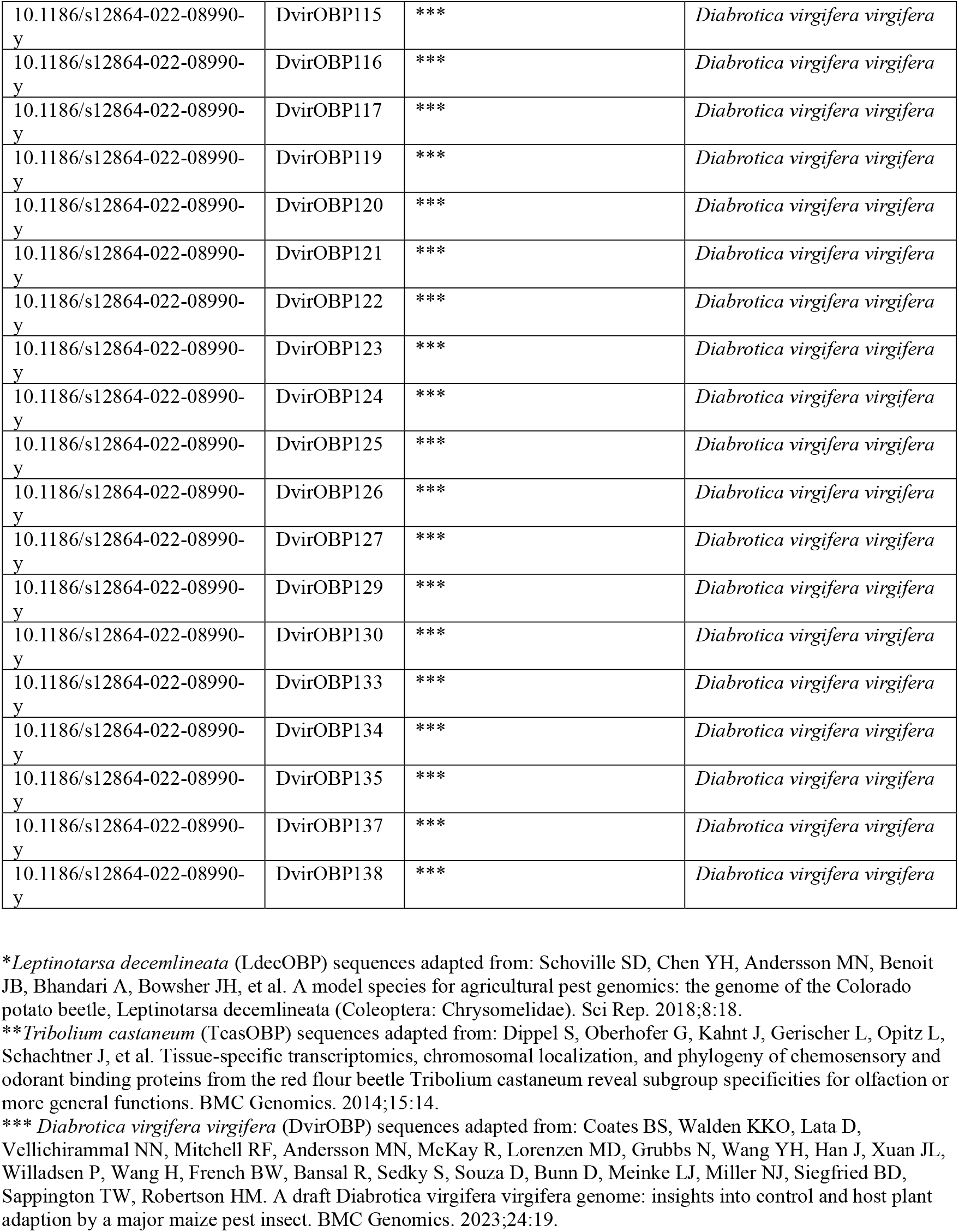
Protein sequences of insect OBP genes are used to construct a maximum likelihood (ML) phylogenetic tree. Sequences not listed with a complementary accession number are adapted from sequences from prior literature that did not feature an accession number.

**Table S2.**
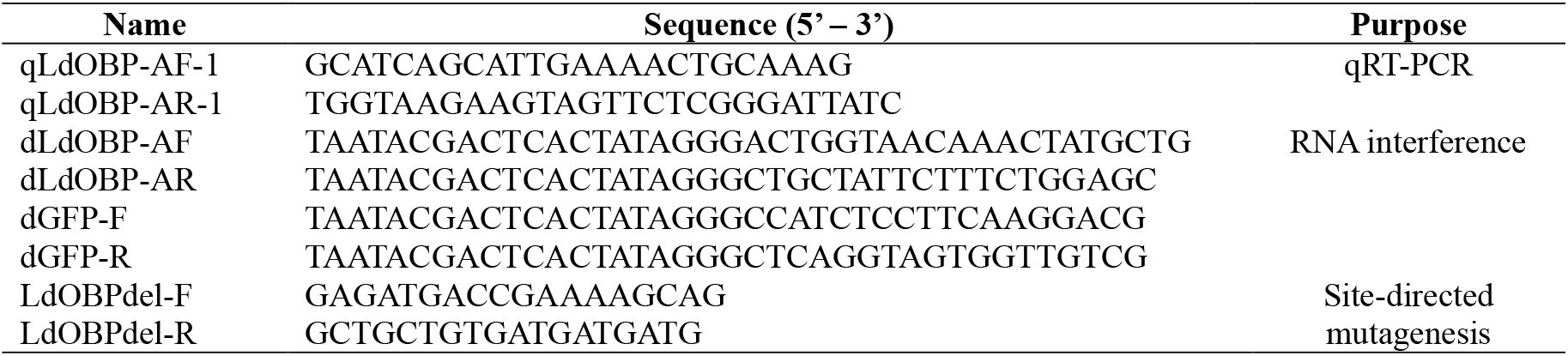
Primers used in this study.

